# Gene copy number is associated with phytochemistry in *Cannabis sativa*

**DOI:** 10.1101/736181

**Authors:** Daniela Vergara, Ezra L. Huscher, Kyle G. Keepers, Robert M. Givens, Christian G. Cizek, Anthony Torres, Reggie Gaudino, Nolan C. Kane

**Affiliations:** Kane Laboratory, Department of Ecology and Evolutionary Biology, University of Colorado Boulder, Boulder, Colorado, USA; Steep Hill Inc., 1005 Parker Street, Berkeley, California, USA

**Keywords:** cannabinoid, CBD, chemotype, copy number variation, hemp, marijuana, metabolic pathway, THC

## Abstract

Gene copy number variation is known to be important in nearly every species where it has been examined. Alterations in gene copy number may provide a fast way of acquiring diversity, allowing rapid adaptation under strong selective pressures, and may also be a key component of standing genetic variation within species. *Cannabis sativa* plants produce a distinguishing set of secondary metabolites, the cannabinoids, many having medicinal utility. Two major cannabinoids -- THCA and CBDA -- are products of a three-step biochemical pathway. Using genomic data for 69 *Cannabis* cultivars from diverse lineages within the species, we found that genes encoding the synthases in this pathway vary in copy number, and that the cannabinoid paralogs may be differentially expressed. We also found that copy number partially explains variation in cannabinoid content levels among *Cannabis* plants.

## Introduction

Gene copy number (CN) varies among individuals of the same species, which may have considerable phenotypic impacts (Stranger et al., 2007). CN variation occurs most commonly via gene duplication (Stranger et al., 2007; Zhang, 2013). Both genome size and complexity can be increased by gene duplication (Zhang, 2013), and new genes can be adaptive (Long, 2013). CN variation seems to be related to gene function, with those encoding biochemical pathway hubs tending to have lower duplicability and evolution rates (Yamada and Bork, 2009). The genes encoding for proteins that interact with the environment reportedly have a higher duplicability (Prachumwat and Li, 2006; Yamada and Bork, 2009), particularly, stress-response genes in multiple plant systems have a high mutation rate (Gaines et al., 2010; Hardigan et al., 2016). Therefore, CN variation can provide a path to rapid evolution in strong selective regimes (Gaines et al., 2010), such as changing environments (Żmieńko et al., 2014; Hardigan et al., 2016) or domestication (Swanson-Wagner et al., 2010; Ollivier et al., 2016).

Three general modes of persistence of duplicated genes that may lead to CN variation have been proposed. The first mode of persistence is concerted evolution, in which the gene copies maintain similar sequence and function but the concentration of the gene product is augmented (Lynch, 2007; Zhang, 2013). The second mode of persistence is neofunctionalization in which a gene copy acquires a novel function (Lynch, 2007; Zhang, 2013). Finally, in subfunctionalization, the original function of the gene becomes split among the copies (Lynch, 2007; Zhang, 2013).

CN variants are often selected during domestication (Swanson-Wagner et al., 2010; Ollivier et al., 2016). Recently, humans have intensively bred for high levels of THCA (delta-9-tetrahydrocannabinolic acid) and CBDA (cannabidiolic acid) (ElSohly et al., 2000; ElSohly and Slade, 2005; Volkow et al., 2014; ElSohly et al., 2016), the two most abundant and well-studied secondary metabolites (also referred to as specialized metabolites) produced by *Cannabis sativa*. This angiosperm from the family Cannabaceae (Bell et al., 2010), produces numerous secondary metabolites called cannabinoids, which are a primary distinguishing characteristic of this plant. These two compounds -- THCA and CBDA -- when heated are converted to the neutral forms −9 tetrahydrocannabinol (THC) and cannabidiol (CBD), respectively (Russo, 2011), which are the forms that interact with the human body (Hart et al., 2001). These compounds have a plethora of both long-known and recently-discovered medicinal (Russo, 2011; Swift et al., 2013; Volkow et al., 2014) and psychoactive properties (ElSohly and Slade, 2005) and are most abundant in the trichomes of female flowers (Sirikantaramas et al., 2005; Gagne et al., 2012). The enzymes responsible for their production, THCA and CBDA synthases (hence THCAS and CBDAS), are alternative end catalysts of a biochemical synthesis pathway (Figure 1; (Sirikantaramas et al., 2005; Gagne et al., 2012; Page and Boubakir, 2014). As *Cannabis*, has had a long history of domestication (Li, 1973; 1974; Russo, 2007), with recent intense selection for THCAS and CBDAS (ElSohly et al., 2000; ElSohly and Slade, 2005; Volkow et al., 2014; ElSohly et al., 2016), CN variation is likely to be found in these synthases (McKernan et al., 2015; Weiblen et al., 2015; Grassa et al., 2018; Laverty et al., 2019). Cannabinoids are thought to abate stresses such as UV light or herbivores (Langenheim, 1994; McPartland et al., 2000; Sirikantaramas et al., 2005), and certain *Cannabis* chemovars contain higher THCA concentrations (e.g. “marijuana-type” cultivars), while other *Cannabis* chemovars contain higher CBDA concentrations (e.g. hemp and high-CBDA “marijuana” varieties) (de Meijer et al., 1992; Rustichelli et al., 1998; Mechtler et al., 2004; Datwyler and Weiblen, 2006).

**Figure 1.**
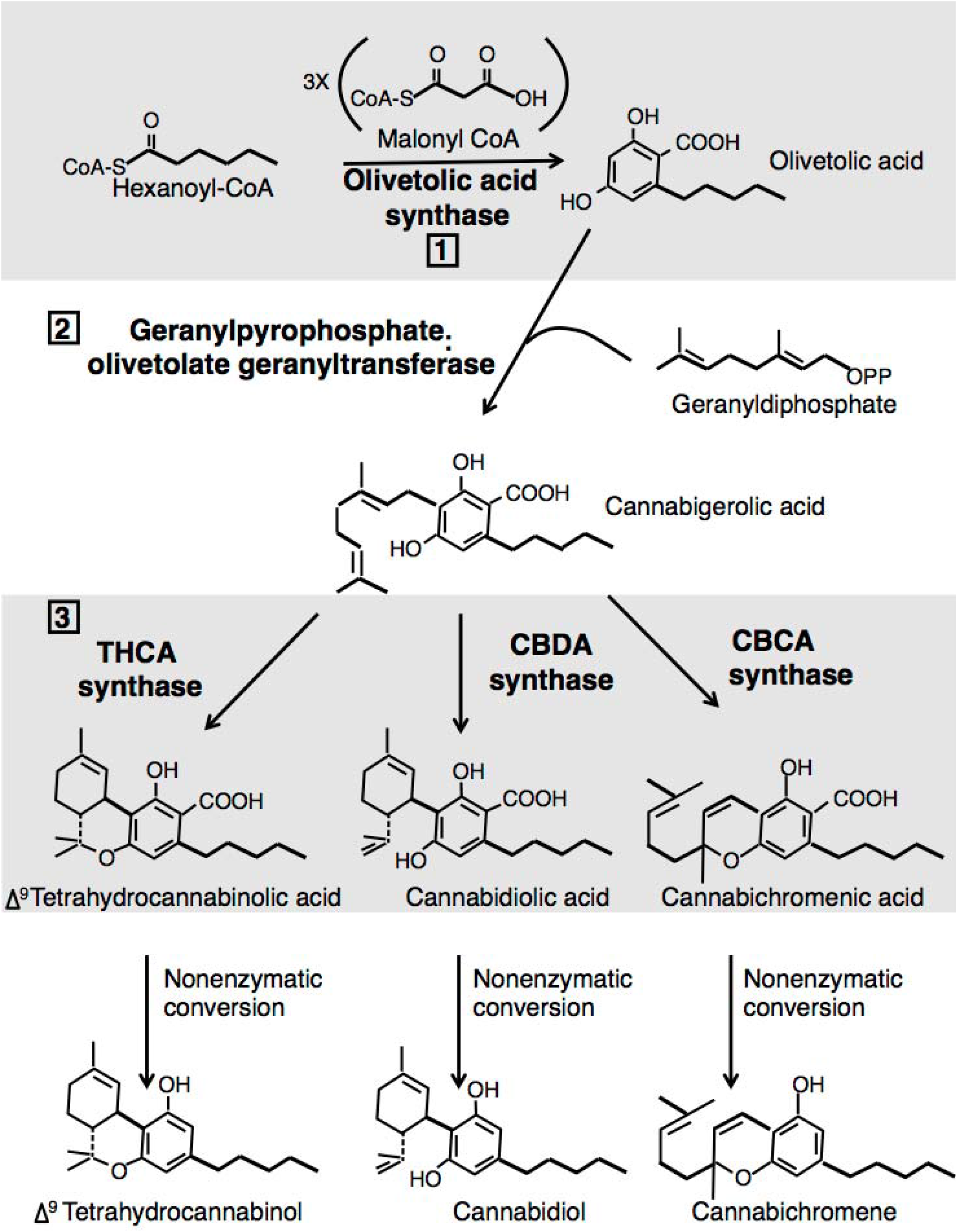
Cannabinoid Synthesis Pathway. The three-step biochemical pathway that produces the medically important cannabinoids in the trichomes of *C. sativa* flowers. Each enzymatic step is labeled with a number: 1) olivetolic acid synthase produces olivetolic acid; 2) olivetolate geranyltransferase produces CBGA; 3) THCA synthase, CBDA synthase, and CBCA synthase produce THCA, CBDA, or CBCA, respectively. The compounds are transformed to their neutral form (THC, CBD, and CBC) with heat in a nonenzymatic conversion. Figure based on Page and Boubakir 2014.

It was thought that allelic variation in the final enzymes in the pathway, THCA and CBDA synthases, determined the predominant cannabinoid composition (de Meijer et al., 1992; de Meijer et al., 2003; Hillig and Mahlberg, 2004; Pacifico et al., 2006; Onofri et al., 2015). However, it has recently been established that there are multiple genes in close proximity that are responsible for the production of cannabinoids (McKernan et al., 2015; Weiblen et al., 2015; Grassa et al., 2018; McKernan et al., 2018; Laverty et al., 2019). Therefore, an alternative explanation for observed phytochemical diversity is that CN variation may contribute to different cannabinoid phenotypes in the *C. sativa* cultivars (McKernan et al., 2015)

Given the medical importance of this pathway and the possibility of CN variation in the genes that encode their enzymes, we explored the inter- and intra-cultivar differences in these genes. Using two de novo *C. sativa* genome assemblies and additional 67 WGS datasets from a diversity of cultivars, we addressed three questions:

**1)** Do lineages differ in number of cannabinoid synthase paralogs? **2)** Does cannabinoid content correlate to the number of respective synthase paralogs by cultivar? **3)** Do cannabinoid synthase paralogs vary in expression level by tissue and cultivar?

## Materials and Methods

### Genome assemblies and gene annotation within the assemblies

We used two different genome assemblies: The first was from a high-THCA marijuana-type male, Pineapple Banana Bubba Kush (PBBK), sequenced using PacBio Single-Molecule Real-Time (SMRT) Long-Read (LR) technology (Eid et al., 2009; Rhoads and Au, 2015), provided by Steep Hill, Inc. (NCBI GenBank WGS accession number MXBD01000000). The second assembly was constructed in 2011 from a high THCA dioecious female marijuana-type Purple Kush (PK) plant, sequenced on the Illumina platform (van Bakel et al., 2011). This was until recently the best *Cannabis* assembly publicly available (Vergara et al., 2016). Most results from this assembly will be given in the Supporting Information. Both assemblies vary in their completeness, as each have some missing BLAST (Altschul et al., 1990; Gish and States, 1993) hits as described below and in the Supporting Information. Each assembly has some duplicated regions, with patterns of coverage suggesting that allelic variation at heterozygous loci lead to two different sequences assembled at a single genomic location. Because both are flawed due to these and other likely misassemblies (Vergara et al., 2016), it was necessary to use both assemblies, which allowed us to find at least one hit for every gene of the pathway in order to understand the whole cannabinoid pathway.

We found two high identity hits containing two exons each to the olivetolic acid synthase gene in the PK assembly (see Supporting Information), and one hit with ten exons to the olivetolate geranyltransferase in the PBBK assembly (see Supporting Information). The two olivetolic acid synthase hits in the PK assembly were found using *C. sativa* OLS olivetol synthase (NCBI accession AB164375.1), and each had a percent identity of more than 80% and an alignment length of at least 1000bp. We found the single hit to the olivetolate geranyltransferase with the mRNA sequence patented by Page and Boubakir (2014) (Page and Boubakir, 2014) -- exclusive to the PBBK assembly -- had a percent identity score of more than 97% (see Supporting Information Tables S1 and S2).

We found 11 and five BLAST hits for putative CBDA/THCA synthase genes in the PBBK and PK assembly, respectively, for a total of 16 potential paralogs in the CBDAS/THCAS gene family (see Supporting Information Table S1). Based on percent-identity scores, we found a hit in each assembly that appears to code for THCAS. We identified two hits in the PBBK and one in the PK assemblies that likely code for CBDAS. We used the CBDAS and THCAS cDNA sequences as reference with NCBI accession numbers AB292682.1 and JQ437488.1, respectively. We also found one hit in the PBBK assembly to the gene producing the third product variant of this pathway, cannabichromenic acid (CBCA) using a cDNA sequence as a reference (Page and Stout, 2017).

We constructed a maximum likelihood (ML) tree using the default parameters in MEGA version 7 (Kumar et al., 2016) with the 16 CBDA/THCA synthase gene family from both assemblies to understand the relationships between them (Figure 2). In order to discern the relationship between the CBDA/THCA synthase gene family, we identified putative homologs of CBDAS/THCAS in closely related species using a tblastx search against NCBI’s non-redundant database. We chose tblastx in lieu of blastx because it allows comparison of nucleotide sequences without the knowledge of any protein translation (Wheeler and Bhagwat, 2007). We included 14 sequences from three species from the order Rosales, two of them also from the family Cannabaceae -- *Trema orientale* and *Parasponia andersonii* with four and three sequences respectively – and a more distantly related species from the family Moraceae as an outgroup, *Morus notabilis*, with seven sequences. Therefore, our ML tree included a total of 30 putative CBDAS/THCAS homolog sequences, 16 from *Cannabis*, seven from two other species in the Cannabaceae, and seven from the outgroup *Morus*. All sequences are deposited on Dryad digital repository (link).

**Figure 2.**
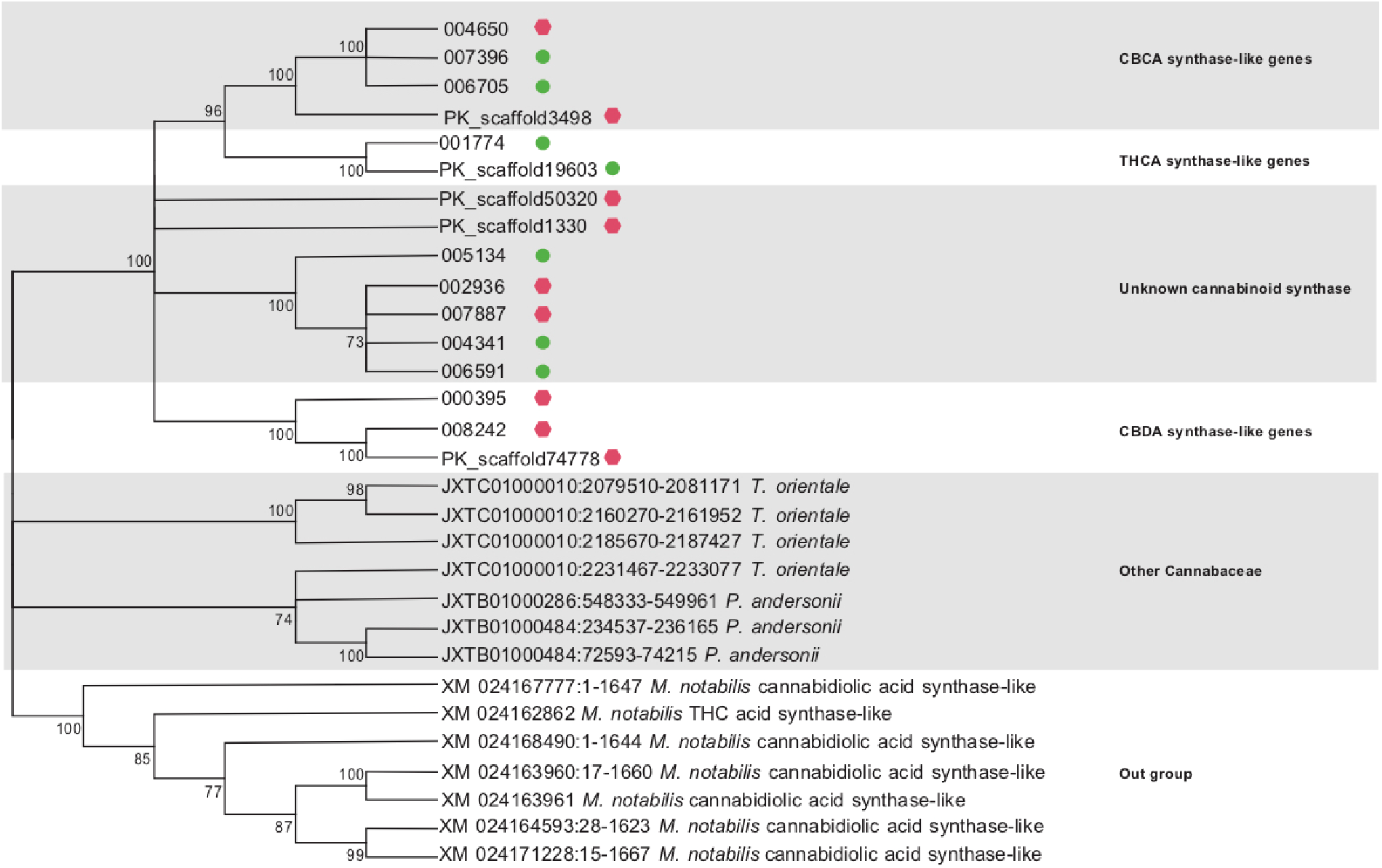
ML Tree with paralogs from the CBDA/THCA synthase family. Relationship between 16 paralogs (11 from the PBBK assembly (prefix “00”) and five from the PK assembly (prefix “PK_scaffold”)). Green circles indicate full-length reading frames, red hexagons indicate truncated reading frames with homology to reference proteins extending beyond stop codons located within them. Paralogs are indicated to be CBDAS-like, THCAS-like or CBCAS-like. Many of the homologs have unknown function. Also included are two other species from the family Cannabaceae, *Parasponia andersonii* and *Trema orientale* with three and four sequences respectively. The outgroup are sequences from the closely related species from the family Moraceae *Morus notabilis*. NCBI accession numbers for each of the proteins listed in the tree. All nucleotide data found on the Dryad repository (XXXX).

Finally, for the 16 sequences we found in the PBBK and PK assemblies, we calculated genetic distance and nucleotide composition using MEGA, and compared the non-synonymous to synonymous sites ratio between sequences with SNAP (Korber, 2000).

### Genomic sequences, alignment, and depth of coverage calculation

We used 67 Illumina platform whole genome shotgun sequence libraries available from various *Cannabis* cultivars (see Supporting Information Table S2) from three major lineages within *C. sativa* (FLOCK; (Duchesne and Turgeon, 2012) groups: Broad Leaf Marijuana-type (broad-leaf), Narrow Leaf Marijuana-type (narrow-leaf), and hemp (Lynch et al., 2016). These genomes have raw read lengths from 100 to 151bp. For detailed information on sequencing and the library prep for these 67 genomes refer to Lynch et al., 2016.

We aligned the 67 libraries to both assemblies using Burrows-Wheeler alignment (BWA) version 0.7.10-r789 (Li and Durbin, 2009), then calculated the depth of coverage using samtools version 1.3.1-36-g613501f (Li et al., 2009). The expected coverage at single copy sites was calculated with the aligned data divided by the genome size (see Supporting Information Table S2), estimated to be 843 Mb for male and 818 Mb for female *Cannabis* plants (van Bakel et al., 2011). Subsequently, the estimated copy number for each cannabinoid sequence was calculated as the average depth across that sequence divided by the expected coverage.

Intrinsic similarity among paralogous genes -- and thus probability that reads from different loci align to the same paralog -- precluded establishing specific SNPs. However, we calculated the number of possible gene paralogs encoding each enzyme in the three terminal steps of CBDA/THCA synthesis (Figure 1) for each cultivar using coverage from both assemblies. The scaled depth was therefore used as a measure of gene CN for each cultivar.

To determine the highest total number of genes per cultivar for CBDAS/THCAS, the depth of coverage was calculated for each library when aligned to the PBBK assembly that had been modified to include only one paralog (PBBK scaffold 001774).

### Gene CN statistics

Differences in the estimated gene CN between the cultivars for each of the 16 CBDAS/THCAS gene family were determined using one-way ANOVAs on the CN of each gene as a function of the lineages (narrow-leaf, broad-leaf, hemp), with a later *post hoc* analysis to establish one-to-one group differences. Three ANOVAs were also performed for each of the lineages to determine within-group variation. The cultivars were then compared with either an ANOVA for cultivars with more than two samples (Carmagnola and Afghan Kush) or a paired t-test for those with two individuals (Chocolope, Kompolti, Feral Nebraska, Durban Poison, and OG Kush; see Supporting Information). Additionally, we performed a Phylogenetic Generalized Least Squares (PGLS) model with the package NLME (Pinheiro et al., 2014) on the R statistical platform (Team, 2013) to determine possible correlations between the depths of each paralog correcting for relatedness between cultivars.

### Phenotypic Analysis

#### Chemotypes

Cannabinoid concentration profiles (chemotypes) were generated by Steep Hill, Inc. following their published protocol (Lynch et al., 2016). Briefly, data collection was performed using high performance liquid chromatography (HPLC) with Agilent (1260 Infinity, Santa Clara, CA) and Shimadzu (Prominence HPLC, Columbia, MD) equipment with 400-6000 mg of sample. We report the estimated total cannabinoid content calculated from the acidic and neutral form of each cannabinoid as in Vergara et al. 2017 and used these values to obtain chemotypic averages for each cultivar. We had the specific chemotypes for eight cultivars which also were sequenced. In these cases, we used individual values instead of the averages (see Supporting Information Table S3).

#### CN vs chemotype correlation

To evaluate the relationship between the estimated gene CN for each of the genes and chemotype, we performed PGLS correlations between the chemotype (phenotype) and the average estimated gene CN per gene (see Supporting Information) while correcting for phylogenetic relatedness. Only cultivars with matching data in the genomic analysis were analyzed, for a total of 35 individuals from 22 different cultivars. The broad-leaf group had 10 individuals from six cultivars, the narrow-leaf had 15 individuals from 13 cultivars, the hemp group had six individuals from one cultivar, and there were four individuals from three cultivars that were not assigned to any group (Lynch et al., 2016). The chemotype data represents 822 individuals from 22 unique cultivars. One caveat of this analysis is that we averaged the chemotypes for most of the shared cultivars except for the eight cultivars for which we had the specific chemotype for that particular genotype (see Supporting Information Table S3). However, an important strength of this average is that effects of environmental variation and statistical noise are minimized, improving our ability to assess genetically-based variation. We also performed PGLS correlations to the sum of all cannabinoids to examine whether CN variation had an effect on overall cannabinoid content.

### Expression Analysis

As a proxy measure of differential expression of the genes on the cannabinoid pathway, we aligned three published RNA sequences derived respectively from the flower and root of Purple Kush (PK) and the flower of the hemp cultivar Finola (van Bakel et al., 2011) to the whole PBBK assembly. We used the Tuxedo suite, which includes Bowtie2 v2.3.4.1 (Langmead and Salzberg, 2012) for RNA alignment, TopHat for mapping v2.1.1 (Trapnell et al., 2009), and Cufflinks v2.2.1 for assembling transcripts and testing for differential expression (Trapnell et al., 2010). We used CummeRbund’s output from the RNA-Seq results (Trapnell et al., 2012).

## Results

### CBDA/THCA synthase family

The quantification of relatedness between the combined 16 CBDA/THCA synthase paralogs drawn from both genome assemblies revealed distinct clusters (Figure 2). Two paralogs, located on contig 001774 and PK scaffold 19603, from the PBBK and PK assemblies respectively, cluster together with 100%-bootstrap support and are related to genes known to be involved in THCA production. Similarly, the paralogs we infer to be CBDA synthases -- two from the PBBK assembly (000395 and 008242) and one from the PK assembly (74778) -- also cluster together. We found a cluster of four genes, three from the PBBK assembly and one from the PK assembly, that we infer to be CBCA synthases. All genes used from the two other Cannabacea species *T. orientale* and *P. andersonii* cluster together. Similarly, the genes from the outgroup *M. notabilis* also form a cluster, to the exclusion of any of the 16 *Cannabis* sequences.

### Gene CN statistics

The one-way ANOVAs for each gene and *post hoc* analysis show that the CN of some of the paralogs differ among the three major cultivar groups (see Supporting Information Table S4 – between-group comparison). However, the *post hoc* analysis with the median from the broad-leaf, narrow-leaf, and hemp groups show that hemps differ from the other two groups in paralog CN, independent of which assembly was used as a reference.

Hemp appears to differ the most from the other two lineages in the copy number of the three CBDAS-like and the two THCAS-like paralogs both between and within lineages (Figure 3), given that for the three paralogs, the hemps have the lowest mean (see Supporting Information Tables S4) and median (Figure 3) CN.

**Figure 3.**
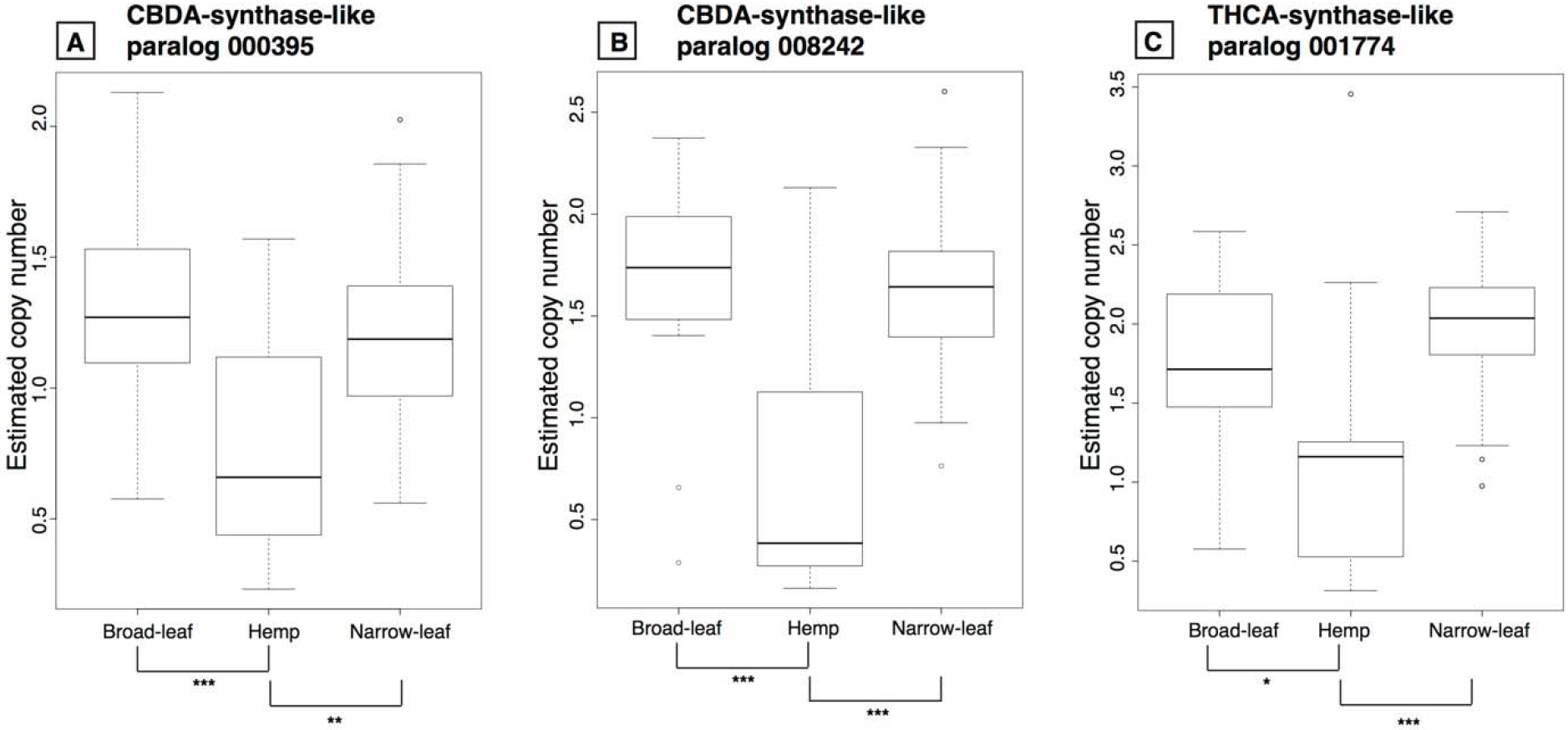
Estimated CN by group for three of the CBDAS/THCAS paralogs. Box plots for three of the paralogs from the 11 total paralogs of the CBDA/THCA synthase family from the PBBK assembly. Panels A and B depict the CBDAS-like genes and panel C is the THCAS-like gene. Significant values between the comparisons are given in the horizontal bars below each panel: *** P<0.001, **P<0.003, *P<0.03. The estimated CN by group from the two CBDAS/THCAS paralogs in the PK assembly are given in Supporting Information figure S1.

The sum of the means of the estimated gene CN per lineage (hemp: μ=15.51; broad-leaf: μ=15.27; narrow-leaf: μ=13.63) is higher than the gene CN on the modified assembly (hemp: μ=11.02; broad-leaf: μ=10.29; narrow-leaf: μ=8.96) with 001774 as the sole representative of its clade (see Supporting Information Table S4). However, the differences between groups in the modified assembly are only marginally significant (F=2.92, p=0.06; Supporting Information Table S4). Despite the only marginally significant differences between groups in the modified assembly, this trend suggests that some of the paralogs have diverged enough that their reads failed to align to the one left in the modified assembly. Still, since some of those genes are truncated, their inclusion in the total CN inflates the sum. Regardless, both estimates show significant variation in CN.

### Phenotypic Analysis

#### CN vs chemotype correlation

After correcting for relatedness, most correlations between the cannabinoid levels and the synthase gene CN lack significance both in the modified and original assemblies (see Supporting Information Table S5). However, the original assemblies had important significant correlations before correcting for relatedness (see Supporting Information Table S5). For CBD chemotypic abundance (after correcting for relatedness) CNs of one (008242) of the two CBDAS-like paralogs significantly but negatively correlate (Figure 4 a,b). Interestingly, the THCAS-like paralog 001774 is also negatively but significantly correlated to CBD accumulation (Figure 4c). For THC chemotypic abundance after correcting for relatedness, all CBDAS/THCAS paralog CNs show significant positive correlations (Figure 5). All other correlations between chemotypic abundance and the multiple gene CNs are given in Supporting Information Table S5. The PGLS correlations to the sum of all cannabinoids behave in a very similar manner as the correlations to single cannabinoids (see Supporting Information Table S5). The patterns shown in figures 4 and 5 are similar to the ones observed when using the PK genome as a reference (see Supporting Information Figure S2 a,b for correlations with percent CBD and Figure S2 c,d for correlations with percent THC).

**Figure 4.**
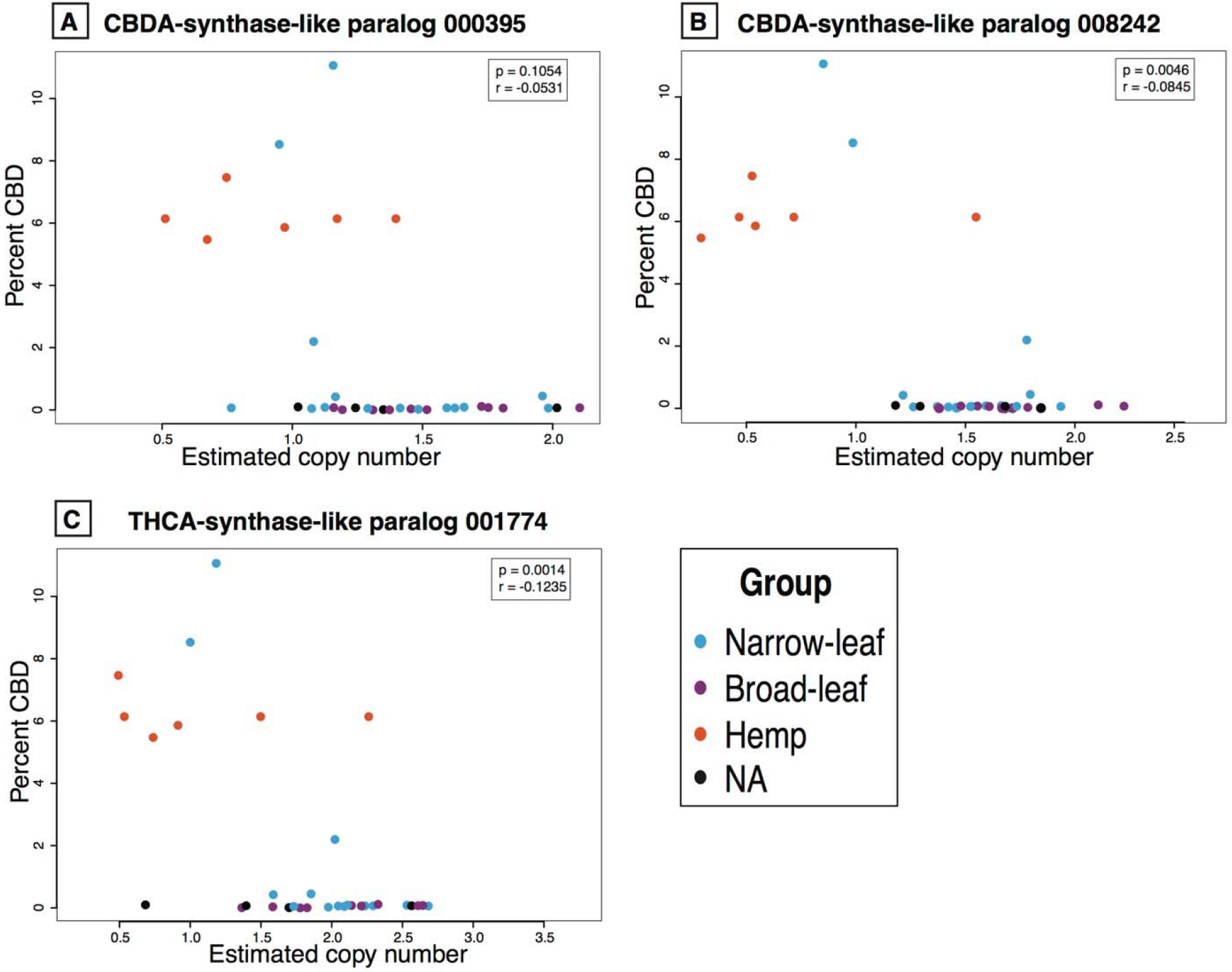
Correlations between the percent CBDA and the estimated CN for the three CBDA/THCA synthase paralogs. Two CBDAS-like genes (panels A and B) and one THCAS-like gene (panel C) correlated to CBDA production. All correlations are negative and those shown in **B** and **C** are significant. Correlation coefficient and p-values in the inset after correction for relatedness. All correlation values between all genes and all cannabinoids are given in Supporting Information Tables S5 and S6, respectively.

**Figure 5.**
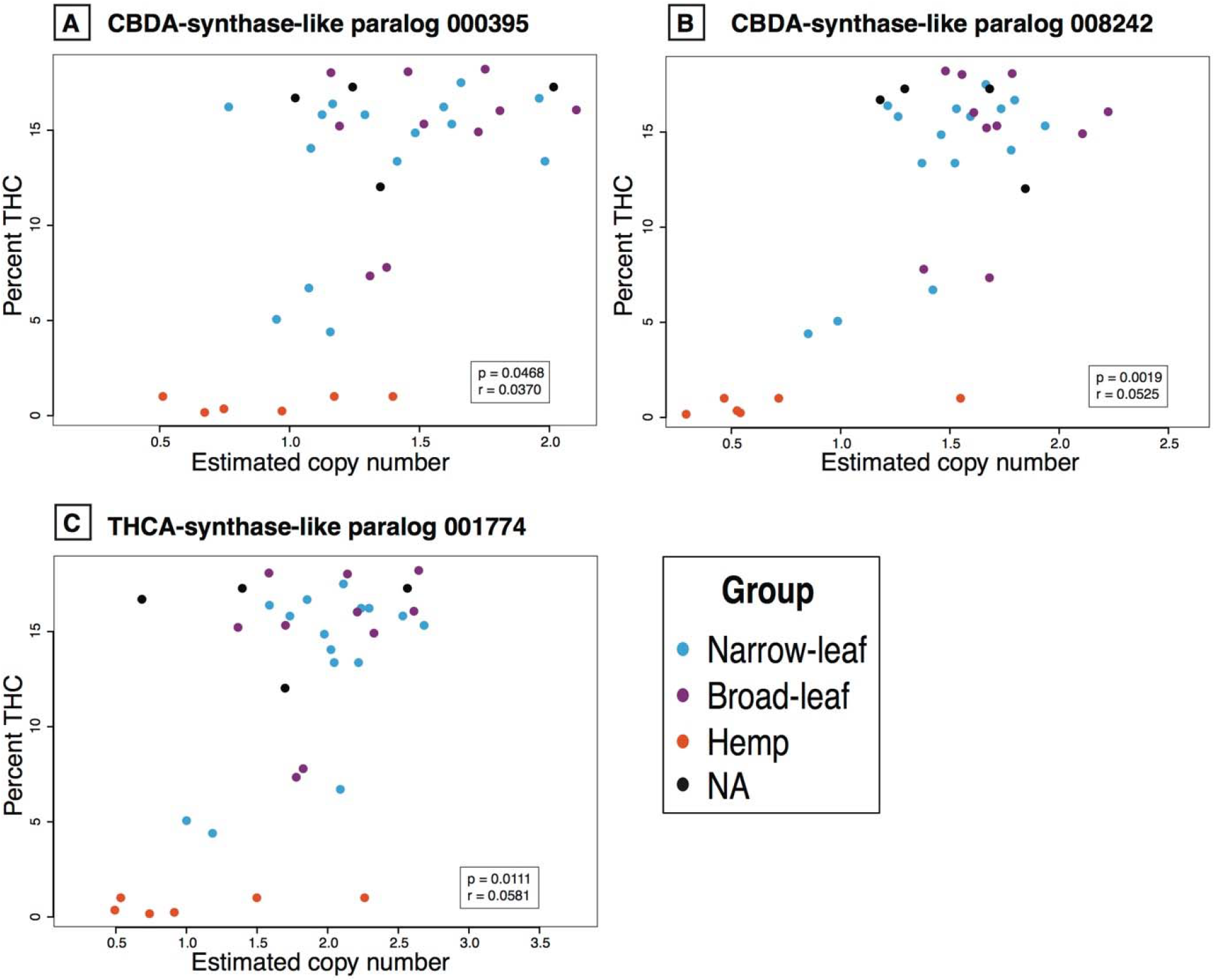
Correlations between the percent THCA and the estimated CN for three CBDA/THCA synthase paralogs. The two CBDAS-like genes (panels A and B) and the one THCAS-like gene (panel C) are positively and significantly correlated at the p<0.05 level to the percent THCA. Correlation coefficient and p-values in the inset after correction for relatedness. All correlation values between all genes and all cannabinoids are given in Supporting Information Tables S5 and S6, respectively.

We found that paralog 006705 had the highest BLAST percent-identity score (99.93%) to the cDNA from the CBCA synthase. Additionally, the two other paralogs that cluster in the same group (007396 and 004650; Figure 1) also show a high-percent identity (99.87% and 99.81% respectively) to CBCA synthase. None of the 16 CBDA/THCA synthase-family paralogs correlate with the accumulation of CBC (see Supporting Information Table S5) after correcting for relatedness. Additionally, the PGLS model with paralogs 007396, 004650, and 006705 did not show any significance. However, three different paralogs (50320, 002936, and 007887) with lower BLAST scores showed a significant correlation with CBC accumulation before correcting for relatedness.

### Expression Analysis

Our proxy expression analysis suggests differences in the gene products between cultivars and tissues (Table 1). Even though the differences are not significant, the marijuana-type cultivar PK seems to express the olivetolate geranyltransferase gene in greater quantities in midflower than Finola the hemp cultivar. This result suggests that the enzymes found upstream of the pathway (such as olivetolate geranyltransferase), may play an important role in the production of cannabinoids, which would be regulated by enzymes found in multiple steps of the pathway. The CBDAS-like paralogs are less abundant in Finola (see Supporting Information Table S3), despite them being significantly more expressed when compared to PK’s mid-flower (Table 1). The THCAS-like paralog is expressed in higher levels in the marijuana-type plant PK, and this comparison is significantly different in the three tissues. The roots of PK seem devoid of transcripts of either the CBDAS or THCAS paralog, likely due to the lack of trichomes in this tissue. These results suggest considerable divergence in expression level, especially given the two order-of-magnitude difference between the expression level of the CBDAS-like paralogs (000395 and 008242) and the THCAS-like paralog (001774).

**Table 1.** Expression for cannabinoid synthase-pathway genes. The expression level for the paralogs related to cannabinoid production vary in both cultivars and tissues. The first column shows each of the paralogs from the PBBK assembly; columns 2,3, and 4 show the average FPKM (fragments per kilobase of transcript per million fragments mapped), which is a measure of expression level proportional to the number of reads sequenced from that transcript after normalizing for transcript’s length, for transcript levels across runs, and for the total yield of the sequencing instrument. Columns 5,6, and 7 show the significance between the pairwise tissue comparison, and finally column 8 shows the group for each of the paralogs.

|  |  |  |  | comparisons |  |  |  |
| --- | --- | --- | --- | --- | --- | --- | --- |
| Paralog | PK<br>midflower<br>(FPKM) | Finola<br>midflower<br>(FPKM) | PK<br>root<br>(FPKM) | PK<br>midflower<br>- Finola<br>midflower | PK<br>midflower<br>- PK root | Finola<br>midflower<br>- PK root | Group |
| 003891 | 243.5 | 16.5 | 0 | NS | P<0.05 | NS | Olivetolate<br>geranyltransferase |
| 006591 | 4.39 | 0.22 | 0 | NS | P<0.01 | NS | Unknown<br>cannabinoid<br>synthases |
| 007887 | 4.383 | 0.221 | 0 | NS | P<0.0001 | NS |  |
| 004341 | 4.38 | 0.22 | 0 | NS | P<0.01 | NS |  |
| 002936 | 4.24 | 0 | 0 | P<0.01 | P<0.01 | NS |  |
| 005134 | 0 | 3.52 | 0 | P<0.01 | P<0.01 | NS |  |
| 000395 | 0.084 | 2.516 | 0 | NS | NS | P<0.01 | CBDAS-like |
| 008242 | 0.468 | 2.75 | 0 | NS | NS | P<0.03 |  |
| 001774 | 484.73 | 1.48 | 0 | P<0.03 | P<0.0001 | P<0.03 | THCAS-like |
| 007396 | 142.91 | 6.08 | 0 | P<0.001 | P<0.0001 | P<0.0001 | CBCAS-like |
| 004650 | 140.94 | 5.67 | 0 | P<0.003 | P<0.0001 | P<0.0001 |  |
| 006705 | 146.99 | 6.05 | 0 | P<0.003 | P<0.0001 | P<0.0001 |  |

## Discussion

In this study, we estimated the CN for the genes encoding enzymes catalyzing three of the main reactions of the biochemical pathway that produces cannabinoids (Figure 1) in the plant *C. sativa*. Although CN variation in some genes involved in cannabinoid production has been previously reported (van Bakel et al., 2011; McKernan et al., 2015), in our study we estimate CN variation in multiple steps of the biochemical pathway in 67 *Cannabis* genomes from multiple varieties within the broad-leaf, narrow-leaf, and hemp groupings using two genome assemblies constructed via complementary technologies.

Our results suggest that synthases for the cannabinoid pathway are highly duplicated and that plants probably use and express the paralogs of these genes differently in specific tissues. Gene CN variation has also been found to be associated with SNP variation and both factors can influence gene expression (Stranger et al., 2007). Our results suggest that this is the case for quantitative and qualitative (amount and type) cannabinoid diversity, which seems to be a product of sequence in agreement to previous research (Onofri et al., 2015), CN variation (McKernan et al., 2015), and expression. The effect of CN variation in relation to these other factors that may affect cannabinoid phenotype is an important topic for further study.

### CBDA/THCA synthase family

The lack of dN/dS value differences and the short genetic distance (see Supporting Information Table S6) suggest that the THCAS/CBDAS gene paralogs arose from a recent duplication event and so have lacked time to accumulate changes. Clusters unique to each of the two assemblies (Figure 2) suggest that either these clades were selectively lost from the opposing assembly or that there exist lineage-specific paralog combinations. The latter would imply that the acquisition and loss of paralogs is rapid enough to show polymorphism at the cultivar level. Interestingly, all three putative CBDAS paralogs from these two high THCA-marijuana-type assemblies bear premature stop codons (Figure 2). This finding supports previous research that suggests that marijuana-type cultivars with high THCA production lack fully functional CBDAS genes (van Bakel et al., 2011; Onofri et al., 2015; Weiblen et al., 2015).

### Gene CN statistics

The difference in CN between hemp and the other two lineages for the three CBDAS-like and the two THCAS-like paralogs (Figure 3) imply that a whole gene cluster was either lost in most of the hemp cultivars or was duplicated in the marijuana-type (broad-leaf and narrow-leaf) individuals. However, even though the hemp group has the lowest mean and median, for many of these genes it has the widest range in gene CN (see Supporting Information Table S4), indicating the widest gene CN variation between the three lineages. CN for these genes differ little between the broad-leaf and narrow-leaf marijuana-types, suggesting similar between-group diversity and higher within-group variation (Figure 3). Our estimates indicate that some of the analyzed individuals from the three different groups could have up to ten copies of CBDAS/THCAS paralogs (see Supporting Information Table S3).

### Phenotypic Analysis

#### CN vs chemotype correlation

There is a positive correlation between accumulation of THC and CN for four of the five paralogs related to CBDA/THCA production, but negative correlation between these paralogs and the accumulation of CBD (Figures 4 and 5, Supporting Information Table S5). This suggests that increasing THCAS gene CN decreases CBDA production possibly due to competition for the mutual precursor, CBGA. Additionally, the THCAS allele from marijuana-type plants appears to be dominant over the THCAS allele from hemp after expression analyses of crossed individuals bearing these alleles, and the CBDAS gene seems to be a better competitor for CBGA even when functional copies of THCAS genes are present (Weiblen et al., 2015). This difference in affinity towards CBGA, and in performance from the various genes and alleles, implies significant contributions from both sequence variation and differences in expression of synthase paralogs to differential accumulation of cannabinoids.

The positive correlation between the CN of the paralogs related to CBDA production (000395 and 008242; Supporting Information Table S7) suggest that these paralogs are physically proximal and were possibly copied in tandem (Weiblen et al., 2015; Grassa et al., 2018). This finding agrees with recent research that found that cannabinoid genes are found in close proximity, in tandem repeats, and surrounded by transposable elements (Grassa et al., 2018; McKernan et al., 2018), which makes sense given that between 43-65% of the *Cannabis* genome consists repetitive sequences (Pisupati et al., 2018). Both paralogs’ CN correlated with the PK paralog 74778 CN (see Supporting Information Table S7), and the three paralogs cluster together (Figure 2), implying that the 74778 paralog in the PK assembly is related to CBDA production. However, the CN of the THCAS-like paralog (001774) is not correlated to the CN from the THCAS-like paralog from the PK assembly (Paralog 19603, Supporting Information Table S7) even though they are closely related (Figure 2). Finally, our BLAST analysis to the two newly published assemblies also show that these cannabinoid genes are in close proximity (Table S9), as reported in their respective publications (Grassa et al., 2018; McKernan et al., 2018).

Another factor that can affect the correlation between synthase gene CN and THCA and CBDA levels is the presence of truncated genes. It has been determined that high-THCA marijuana cultivars possess a truncated version of the CBDA synthase (van Bakel et al., 2011; Onofri et al., 2015; Weiblen et al., 2015). The presence of the truncated CBDAS paralogs can explain some of the points in Figure 4 in the bottom right corner where, even though the estimated CN is high (high value on the X axis), the amount of CBD produced is low (low value on the Y axis) due to the premature termination and inability to produce the protein. Truncated genes have also been reported for THCA synthases (van Bakel et al., 2011; Onofri et al., 2015; Weiblen et al., 2015), however we do not see many samples in the bottom right corner with high CN and low THC production (Figure 5).

It is interesting that the individual hemp-type plants have the lowest mean and median CN for the three CBDAS/THCAS paralogs (Figure 3 and Supporting Information Table S4). We expected hemp types to have a higher mean CN of the two paralogs related to CBDA production, given their higher production of CBDA compared to marijuana types (de Meijer et al., 1992; Rustichelli et al., 1998; Mechtler et al., 2004; Datwyler and Weiblen, 2006). However, hemp individuals have a higher mean for other paralogs from the CBDA/THCA synthase family (see Supporting Information Table S4) such as paralog 005134 which has a negative correlation with the production of THCA but positive for CBDA (see Supporting Information Table S5). Finally, recent research suggest that CBDA-dominant lineages seem to produce minor cannabinoids absent in certain THCA lineages, implying the loss of cannabinoid genes in these highly hybridized THCA-dominant cultivars (Mudge et al., 2018). Perhaps these paralogs found in the hemp lineages may be related to these minor cannabinoids.

### Expression Analysis

Variation in expression profiles of the THCAS and CBDAS gene paralogs (Table 1) could be another major contributor to measured phenotypic differences among *Cannabis* cultivars, as seen for genes related to stress response in maize (Waters et al., 2017). This effect may be augmented by the fact that chemotype assays are generally performed on mature flower masses. Variation in transcription is seen for many of the CBDAS/THCAS paralogs by both tissue and cultivar, suggesting differential use of pathway genes. On the other hand, transcripts from most cannabinoid synthase paralog clades are transcribed in greater quantities by the marijuana cultivar PK in marked contrast to the hemp cultivar Finola (Table 1), implying that marijuana cultivars express more diversity in cannabinoid synthase genes. Finally, CN variation can correlate positively or negatively with gene expression (Stranger et al., 2007), which could be the case for THCAS and CBDAS, as may be the particular case for paralog 008242 that has a significant negative correlation with CBDA production.

### CN variation and the cannabinoid pathway

In other plant species such as potatoes and maize, species-specific secondary metabolites accumulating in glandular trichomes confer resistance to pests and the corresponding synthase genes are found in high copy numbers (Hardigan et al., 2016; Waters et al., 2017). This appears to be the case in *Cannabis*. Cannabinoid synthesis appears to be genus-specific and accumulation of cannabinoids in glandular trichomes could be stress-related (Langenheim, 1994; Sirikantaramas et al., 2005). Our results suggest that the CBDA/THCA synthase family has recently undergone an expansion. Previous studies have assumed that CBDAS was the ancestral gene and that THCAS arose after duplication and divergence (Onofri et al., 2015), but since no other species is known to share this biosynthetic pathway it’s not possible to conclusively identify the ancestral state. Our phylogenetic analysis suggests that these cannabinoid genes are specific to *Cannabis*, but in order to conclusively determine which is the ancestral state other closely related extinct and extant species (ie. <u>*Humulus*) remain to be analyzed for the presence of genes related to the</u> CBDA/THCA synthase family.

Regardless, duplication and neofunctionalization of ancestral synthase genes is a likely contributor to chemotype variability. CN variants can serve as a mechanism for species-specific expansion in gene families involved in plant stress pathways (Hardigan et al., 2016; Waters et al., 2017). Additionally, CN variation has been reported in gene families involved in stress response and local adaptation in plants (Hardigan et al., 2016; Waters et al., 2017), and other organisms (Van de Peer et al., 2017), perhaps explaining why all genes in the cannabinoid pathway have been highly duplicated.

The high numbers of paralogs in the CBDAS/THCAS family supports the notion that biosynthesis proteins that have fewer internal metabolic pathway connections have a higher potential for gene duplicability (Prachumwat and Li, 2006; Yamada and Bork, 2009). However, despite both olivetolic acid synthase and olivetolate geranyltransferase operating near the pathway hub, the respective estimated CNs of their paralogs are similar to the CN of CBDA/THCA synthase paralogs (Figure 1, Supporting Information Table S4). Sequence similarity and physical proximity of extant paralogs in the genome (Weiblen et al., 2015; Grassa et al., 2018) promotes tandem duplication, again facilitating rapid expansion of the CBDA/THCA synthase family. Human selection since the ancient domestication of this plant has likely played a role, as it did with CN in resistance genes in the plant *Amaranthus palmeri* (Gaines et al., 2010) and in the starch digestion gene *Amy2B* during dog domestication (Ollivier et al., 2016). Finally, gene CN variation has been associated with SNP variation and both factors can influence phenotype expression (Stranger et al., 2007).

Our study provides another example of the high association between the CBDA/THCA synthase gene family, which has a very particular relationship, compete for the same precursor molecule (Page and Boubakir, 2014; Page and Stout, 2017), are similar to each other in their chemical structure (Brenneisen, 2007; Flores-Sanchez and Verpoorte, 2008) in their genetic sequence (Onofri et al., 2015), and may exemplify “sloppy” enzymes (Auldridge et al., 2006; Franco, 2011; Chakraborty et al., 2013). These “sloppy” enzymes could convert similar substrates (such as CBGA) into a range of slightly different products, such as CBDA, THCA, or CBCA (Jones et al., 1991).

### Caveats

In addition to the factors previously examined as contributing to the high intrinsic genomic complexity of cannabinoid synthesis pathway regulation, the possible misassembly of both genomes may further confound attempts at precise correlations. The PK contigs have been misassembled probably due to very short reads combined with high heterozygosity. This misassembly in the PK genome may be the reason why, even in further assembly attempts, the cannabinoid synthases are found in different locations in the genome (Laverty et al., 2019) despite other assemblies finding these genes in close proximity (Grassa et al., 2018; McKernan et al., 2018). The scaffolds in the PBBK assembly where the CBDAS/THCAS family genes are located lack sequence similarity beyond the gene borders indicating that these scaffolds likely have not been affected. However, the very similar paralogs that cluster together in the ML tree (Figure 2) could be different alleles of the same gene that were assembled in different scaffolds. Additionally, the finding of some synthases exclusively in one or the other assembly suggests data gaps in both genomes, although the differences may represent true biological variation given the high amount of CN variation among the different *Cannabis* varieties. This second hypothesis, suggesting that these differences are true biological variation is supported by other research (McKernan et al., 2015), and by our BLAST analysis (Table S9) to two newer assemblies.

### Conclusions

In conclusion, returning to our three initial questions: **1)** Do lineages differ in number of cannabinoid synthase paralogs? We found that the measured copy-number of these genes did vary, within and between lineages and possibly within named cultivars given by the differences in CN (see Supporting Information Table S4). **2)** Does cannabinoid content correlate with the number of respective synthase paralogs by cultivar? We found a positive correlation between the accumulation of specific cannabinoids and the CN of certain synthase paralogs. THCA levels are significantly and positively correlated with the CN of several of these paralogs (Figures 4 and 5, and Supporting Information Table S5). Furthermore, the broad-leaf and the narrow-leaf marijuana types each have a higher mean and median for the CN’s of genes related to the production of both THCA and CBDA relative to hemp cultivars. However, CBDA levels are negatively correlated with most of the paralogs related to its production, and the hemp cultivars paradoxically exhibit higher CNs for the PK contig 19603 THCAS-like paralog than for CBDAS paralogs (See Supporting Information Figure S1, Table S5). We found both positive and negative correlations between the production of the other cannabinoids and the CN of some of the paralogs, making it difficult to associate particular cannabinoids with specific paralogs (Figures 3, 4, and Supporting Information Figure S2). **3)** Do cannabinoid synthase paralogs vary in expression level by tissue and cultivar? We observed differential transcription levels of these genes by tissue in conjunction with cultivar (Table 1) which likely adds to the high complexity of correlating paralog CNs with cannabinoid accumulation.

Finally, our findings motivate a pair of general breeding strategies. To boost production of THCA, select parents with higher CNs of THCAS paralogs, whereas for cultivars with more CBDA, select parents with fewer such paralogs. Given that cultivars express synthases from multiple points in the pathway differently (Table 1), all of these genes should be considered for breeding purposes. For exclusive production of either THCA or CBDA, cross cultivars bearing only truncated paralogs of the opposing synthase genes.

## Supporting information

Supplemental materials and results

## Author Contributions

D.V. analyzed the copy number data, wrote the first draft of the manuscript, conceived and lead the project; E.L.H. wrote bioinformatic pipelines for the depth, GLS models, and expression analyses, organized the manuscript’s code and made it publicly available; K.G.K. helped with the original bioinformatic code and project conception; R.G., A.T., C.G.C.,. designed, supervised, and provided chemotype data collection; R.M.G, AT selected and extracted SMRT-LR template DNA; N.C.K. conceived and directed the project. All authors contributed to statistical analysis and manuscript preparation.

## Acknowledgments

D.V. is the founder and president of the non-profit organization Agricultural Genomics Foundation, and the sole owner of CGRI, LLC. R.G., A.T., C.G.C., and R.M.G. are employees of Steep Hill, Inc. N.C.K. is a board member of the non-profit organization Agricultural Genomics Foundation.

We thank B. Holmes of Centennial Seeds; D. Liles, C. Casad, A. Ledden and J. Cole of The Farm; MMJ America, Medicinal Genomics, A. Rheingold and M. Rheingold of Headquarters; D. Salama, Nico Escondido, Sunrise Genetics, and B. Sievers for providing DNA samples or sequence information; O. Vergara for help with the original bioinformatic code; A. Wiens and Z. Mullen from LISA for help with the PGLS models; and C. Pauli, A. Goebl and the Kane, Flaxman, Safran, and Taylor labs for comments on the manuscript.

**Supporting Information Table S1. Genes from the Cannabinoid Pathway.** Information on the different paralogs from the three-step biochemical pathway, including the gene (column 1), the assembly used for each gene (column 2), the scaffold in which each paralog was found (column 3), the beginning and end positions of each gene within its scaffold (columns 4 and 5), the number of exons (column 6), and the BLAST percent identity (column 7). For the start and end positions (columns 4 and 5), if the gene is found in the reverse strand, it will have a higher value for the start than for the end. For the last column (7) we calculated the identity using the mRNA with accession number AB164375.1 for olivetolic acid synthase, the mRNA sequence patented by Page and Boubakir (2014) for olivetolate geranyltransferase, and the mRNA sequence from THCA with the NCBI accession number JQ437488.1 for both THCA and CBDA. We calculated the average percent identity for the exons from olivetolic acid synthase and olivetolate geranyltransferase but not for THCA and CBDA since they only contain one exon.

**Supporting Information Table S2. WGS information.** Information on each of the 67 WGS sequenced on the Illumina platform used for the depth analysis. Each cultivar has a unique sample ID (column 1), name (column 2), colloquial classification (column 3), lineage (flock group; column 4) determined by (Lynch et al. 2017), ID with the NCBI submission (column 5), the size of the alignment determined with the sam file (column 6), and the scaled depth (column 7) which was determined by dividing the size of the alignment by the genome size.

**Supporting Information Table S3. Average depth and chemotypes.** Columns 1 – 24 show the average depth by cultivar for each of the 19 paralogs analyzed (two for olivetolic acid synthase, one for olivetolate geranyltransferase, and 16 from CBDAS/THCAS), and the paralogs from the modified assemblies (paralog 16618 from the PK assembly, and paralog 001774 from the PBBK assembly for CBDAS/THCAS). Columns 25-29 show five chemotypes for 22 cultivars. The final column indicates whether the chemotype information is an average (Y), a specific value (N), or is absent (0).

**Supporting Information Table S4. Statistics for differences in CN between and within groups, and within repeated strains including modified assemblies.** Statistical results from the ANOVAs (between and within groups, and for the two cultivars that had more than two independent samples each -- Carmagnola and Afghan Kush -- and t-tests (for the cultivars that had only two individuals – Chocolope, Kompolti, Feral Nebraska, Durban Poison and OG Kush). The p-values that are not shown are significant at the p<0.001 level. Calculations in the bottom of the table show the sum of the means for the olivetolic acid synthase paralogs (15717 and 16618) and for the 11 CBDA/THCA synthase paralogs from the PBBK assembly by lineage.

**Supporting Information Table S5. Correlations between the estimated CN of the 19 different paralogs (including the paralogs from the modified assemblies) and the chemotype for five cannabinoids corrected for relatedness.** Column 1 is the gene for each of the enzymes in the pathway, the assembly used for each gene is found in column 2, and the scaffold in which each paralog was found in column 3. None of the estimated CN of any paralog is significant after Bonferroni corrections for multiple comparisons. Entries with an asterisk (*) are values that were significant before correcting for relatedness. The final two columns are the statistics of the correlations between the estimated CN and the sum of all cannabinoids.

**Supporting Information Table S6. Genetic Distance (upper half) and dN/dS ratio (bottom half) for the 16 CBDAS/THCAS paralogs.** The first 11 rows and columns belong to the 11 paralogs from the PBBK assembly; the remaining five rows and columns correspond to the five paralogs from the PK assembly. Each entry is the pairwise comparison between two paralogs for either the genetic distance (upper half) or the dN/dS ratio (bottom half).

**Supporting Information Table S7. Correlations between the estimated CN of the 19 different paralogs including the paralogs from the modified assemblies corrected for relatedness.** The estimated CN of some of the paralogs correlate between them, independent of what gene they codify. Bold entries signify values that remain significant after Bonferroni corrections for multiple comparisons, and entries with an asterisk (*) are values that were significant before correcting for relatedness.

**Supporting Information Table S8. Exons and Introns for olivetolic acid and olivetolate geranyltransferase synthases.** Positions of the exons and introns for the two olivetolic acid synthase paralogs from the PK assembly, and the one olivetolate geranyltransferase gene in the PBBK assembly.

**Supporting Information Table S9. BLAST results to two newly published assemblies** in non-peer reviewed archives from Grassa et al., 2018 and McKernan et al., 2018 (Column 1) with 12 and 27 hits (Column 2) respectively, each with a percent identity of more than 80% and a length of more than 1000bp. The 12 hits for Grassa et al.’s assembly are all found in chromosome nine (Column 3), while the 27 hits in McKernan et. al.’s assembly are found in seven different unplaced scaffolds (Column 3). We used the THCAS and CBDAS with NCBI accession numbers JQ437488.1 and AB292682.1 respectively. For CBCAS we used the sequence from Page and Stout, 2017. All hits for these three synthases have the same starting position (Column 4) but different ending positions, percent identity, and alignment length, for THCAS (Columns 5-7), CBDAS (Columns 8-10), and CBCAS (Columns 12-14), respectively. The paralogs are ordered according to their percent identity to THCAS, though the order for their resemblance to CBDAS and CBCAS are reported as well, in columns 11 and 15, respectively. The hit with the highest percent identity to each of the synthases in both assemblies is bolded.

**Supporting Information Figure S1. Estimated CN by group for the two of the CBDAS/THCAS paralogs from the PK assembly.** Box plots for two of the paralogs from the 5 total paralogs of the CBDA/THCA synthase family from the PK assembly. Panel A is CBDAS-like gene and panel B is the THCAS-like gene. Significant values between the comparisons are given in the horizontal bars below each panel: *** P<0.001, **P<0.003, *P<0.03.

**Supporting Information Figure S2. Correlations between the percent CBDA and the percent THCA and the estimated CN for two CBDA/THCA synthase paralogs from the PK assembly.** The percent CBDA (Panels A and B) is negatively correlated -- while the percent THCA (Panels C and D) is positively correlated -- with CNs of both CBDAS-like paralog 74778 and THCAS-like paralog 19603 from the PK assembly. Correlation coefficients and p-values in the inset after correction for relatedness. Only the CBDAS-like paralog 74778 is significantly correlated with both CBDA (Panel A) and THCA (Panel C), while the THCAS-like paralog 19603 lacks significance (Panels B and D). All correlation values between all genes and all cannabinoids are given in Supporting Information Table S6.

