## Supplemental materials and results for "Gene copy number is associated with phytochemistry in *Cannabis sativa*"

**Methods**

Genomic assemblies and gene determination within the assemblies

Using nucleotide BLAST against the complete mRNA sequences of the synthases in the cannabinoid pathway, we identify the loci encoding these enzymes and their positions in both assemblies. We found two high-homology hits containing two exons each to the olivetolic acid synthase gene in the PK assembly, one hit with ten exons to the olivetolate geranyltransferase in the PBBK assembly, and 11 and five hits to the CBDA/THCA synthases in both the PBBK and the PK assemblies respectively (Tables S1 and S8).

The two olivetolic acid synthase hits in the PK assembly were found using *C. sativa* OLS olivetol synthase (NCBI accession AB164375.1), and each had a percent identity of more than 80% and an alignment length of greater than 1000bp. We did not find sequences with these criteria in the PBBK assembly. We compared the pairwise genetic distance between the exons and the introns of the duplicate sequences using the program MEGA version 7 (Kumar et al., 2016), and the non-synonymous to synonymous sites ratio between exons with the program SNAP (Korber, 2000).

Our single hit to the olivetolate geranyltransferase found with the mRNA sequence patented by Page and Boubakir (2014) (Page and Boubakir, 2014) -exclusive to the PBBK assembly- had a percent identity score of more than 97% (Tables S1 and S2). We only observed partial-length hits in PK assembly, which could be pseudogenes. Given that olivetolate geranyltransferase is an important enzyme in this pathway, it’s likely that the PK assembly due to missasembly, lacks the full-length gene.

The THCA synthase coding sequence (NCBI accession JQ437488.1) yielded 11 qualifying hits in the PBBK assembly and five in the PK assembly with a percent identity of more than 80% and an alignment length of more than 1100bp, for a total of 16 paralogs in the CBDA/THCA synthase family from both assemblies. Each complete THCA and CBDA genes is comprised of one exon of approximately 1600bp each. With a recently published cDNA from CBCA synthase, we found that paralog 006705 had the highest BLAST percent-identity score (Page and Stout, 2017). We did an additional BLAST analysis to two newly published assemblies (Grassa et al., 2018; McKernan et al., 2018) to corroborate whether the CBDA/THCA synthase gene family were found in higher copies on more complete assemblies.

Genomic sequences, alignment, and CN calculation

In order to calculate the approximate gene copy number in every individual sample, we first generated 134 files (one for each of 67 cultivars aligned to each assembly) containing the alignment depth per position for each base pair. We estimated the expected sequencing depth of the single-copy portions of each of the 67 samples from the medians of histograms generated from the alignment depths of each position in the genome. The expected gene copy number in each sample was then calculated as the average alignment depth within the coding sequence of each gene normalized by the expected depth in the single-copy portion of the respective assembly from which the reference synthase belonged.

We compared CNs between lineages (FLOCK groups) by computing the median for each of the 19 genes (two olivetolic acid synthase, one olivetolate geranyltransferase, 16 CBDA/TCHA synthases) that were used in the one-way ANOVAs. For cultivars containing more than two sequenced individuals (Carmagnola and Afghan Kush), we compared CN estimates via depth in every position of each gene with one-way ANOVAs. We compared the cultivars limited to two sequenced individuals (Chocolope, Kompolti, Feral Nebraska, Durban Poison, and OG Kush) with t-tests.

Similar to the analysis applied to the CBDA/THCA synthases, the highest total number of genes per cultivar for olivetolic acid synthase was determined using depth of coverage calculated for each library when aligned to the PK assembly that had been modified to include only one paralog of (paralog 16618).

Gene CN statistics

As described for the 16 CBDA/THCA genes, in order to estimate the differences in gene CN between the cultivars for each of the two olivetolic acid synthase and one olivetolate geranyltransferase, we performed one-way ANOVAs on the CN of each gene as a function of the FLOCK lineages (narrow-leaf, broad-leaf, hemp). We established one-to-one group differences with a posterior posthoc analysis. As with the CBDA/THCA synthase family, we performed three ANOVAs for each lineage to determine within-group variation, and then compared the cultivars sampled more than twice (Carmagnola and Afghan Kush) with an ANOVA or with a paired t-test for those represented by only two individuals (Chocolope, Kompolti, Feral Nebraska, Durban Poison, and OG Kush). Finally, we performed separate ANOVAs on the exons and introns for the modified olivetolic acid synthase alignment.

We used Generalized Least Squares (GLS) models from the NLME package on the R statistical framework to correlate the CN between each of the paralogs (Table S7) correcting for cultivar relatedness calculated by Lynch et al. (2017). This correction allows us to better understand whether the patterns are produced because of CN variation independent of whether the individuals are related to each other. In other words we are controlling for possible kinship.

Phenotypic Analysis

*Correlation between read alignment depth and chemotype*

We used GLS to establish whether the amount of each cannabinoid was explained by CNs of the putative synthase genes (Table S6), correcting for relatedness. Additionally, we tested whether the level of CBD as a response variable was explained by the three CBDA-like synthase paralogs from both assemblies (000395 and 008242 from the PBBK assembly and 74778 from the PK assembly; Figure 2) in a first model, or to the paralogs that had a significant one-to-one correlation with CBD (001774 and 008242 from the PBBK assembly, and 74778, and 50320 from the PK assembly; Figure 2, Table S7) in a second model.

Similarly, we ran two GLS models with THC level as the response variable. In the first model we tested for a relationship between the two THCA-like paralogs 19603 from the PK assembly and 001774 from the PBBK assembly (Figure 2). In the second model we used the paralogs that showed a significant correlation with THC (001774, 000395, and 008242, from the PBBK assembly and 74778 from the PK assembly; Figure 2, Table S7) in a second model.

Finally, for CBC as a response variable, we implemented one GLS model to test for a relationship between the paralogs possibly responsible for its production all from the PBBK assembly (006705, 007396, and 004650, Figure 2).

All of these models were corrected for relatedness between cultivars using relatedness values obtained from Lynch et al (2017).

Expression Analysis

We used the Tuxedo suite to understand the differential expression of the genes in the cannabinoid pathway. The length in base pairs for the olivetolate geranyltransferase and the 11 CBDA/THCA paralogs found with BLAST in the PBBK assembly were input into TopHat (Trapnell et al., 2010; Trapnell et al., 2012) as a standard gene transfer format (GTF) file. We were able to measure the differential expression of these paralogs by aligning three published RNA sequences from the flower and root of Purple Kush (PK) and the flower of the hemp cultivar Finola^24^.

**Results**

*Olivetolic acid synthase*

The two previously reported (van Bakel et al., 2011) paralogs for olivetolic acid synthase in the PK assembly have pairwise genetic distances of 0.007 and 0.059 between the exons and intron, respectively, representing an eight-fold faster mutation rate in the selectively-neutral intron. The dN/dS ratio between the exons is 0.27. The accelerated mutation rate of the intron combined with a dN/dS < 1 support the conclusion that both copies of the olivetolic acid synthase are under purifying selection. Additionally, the similarity of intron position and length (Tables S1 and S8) implies that the whole DNA portion including both exons and introns was duplicated.

Despite their similarity in sequence, the paralogs behave distinctly as the correlations between CN (Table S7) and chemotype (Table S6) differ between them. This difference in behavior between the two paralogs could be due to mutations affecting their catalytic site or impacting their activity at the transcriptional and translational levels.

The modified alignment that included only one of the paralogs for olivetolic acid synthase (16618 from the PK assembly), yielded CN means (hemp: μ=2.56; broad-leaf: μ=3.13; narrow-leaf: μ=2.69) that are very similar to the sum of the means of the estimated CN (hemp: μ=2.53; broad-leaf: μ=3.09; narrow-leaf: μ=2.64). The fact that the mean of the modified assembly and the sum of the CN means are similar suggests that both paralogs are alike and both of their reads align to the modified assembly. The estimated CN for the two exons and intron for olivetolic acid synthase differ, where the CN for the intron is lower than both exons in the three lineages and within the repeated cultivars (Table S5), indicating that fewer reads align to the intron than to the exons. This lower coverage in the intron is likely due to the higher rate of divergence in the intron.

Olivetolate geranyltransferase

We expected a significant positive correlation between the estimated gene CN for olivetolate geranyltransferase (paralog 003891) from the PBBK assembly and the abundance of CBG (Table S7) because this paralog is the closest hit to a vetted CBGA synthase (Table S1)(Page and Boubakir, 2014). The lack of correlation suggests that there might be other olivetolate geranyltransferase paralogs that were missed in both assemblies and that we found a non-functional or re-purposed paralog, or that CBG abundance depends on limiting precursor pools or on conversion rates by the THCA/CBDA/CBCA synthases present.

The CN of the olivetolate geranyltransferase paralog (003891) is negatively correlated with the CN of all of the genes in the CBDA/THCA family (Table S7). In other words, the more olivetolate geranyltransferase gene copies a cultivar has, the fewer gene copies of most of the CBDA/THCA paralogs it possesses.

The enzyme-reaction-rate relationship in metabolic pathways can be complex because one enzyme may catalyze different reactions and multiple enzymes may catalyze the same reaction(Ma and Zeng, 2004). This lack of a one-to-one correlation might explain the negative correlations between the CNs of some of the paralogs and between the gene CN and the production of cannabinoids.

*CBDA/THCA synthase family*

Using the CBDA-like (74778) and the THCA-like (19603) paralogs from the PK assembly, we found slightly different results compared to the paralogs from the PBBK assembly (Figure 3). Hemp appears to differ the most from the other two lineages in the copy number of the CBDA-like genes (Figure S1 panel A) for both assemblies, but not for the THCA synthase paralog from the PK assembly (Figure S1 panel B). Still, the hemp group has the widest range in gene CN (Figures 2 and S1), indicating the widest gene CN variation between the three lineages.

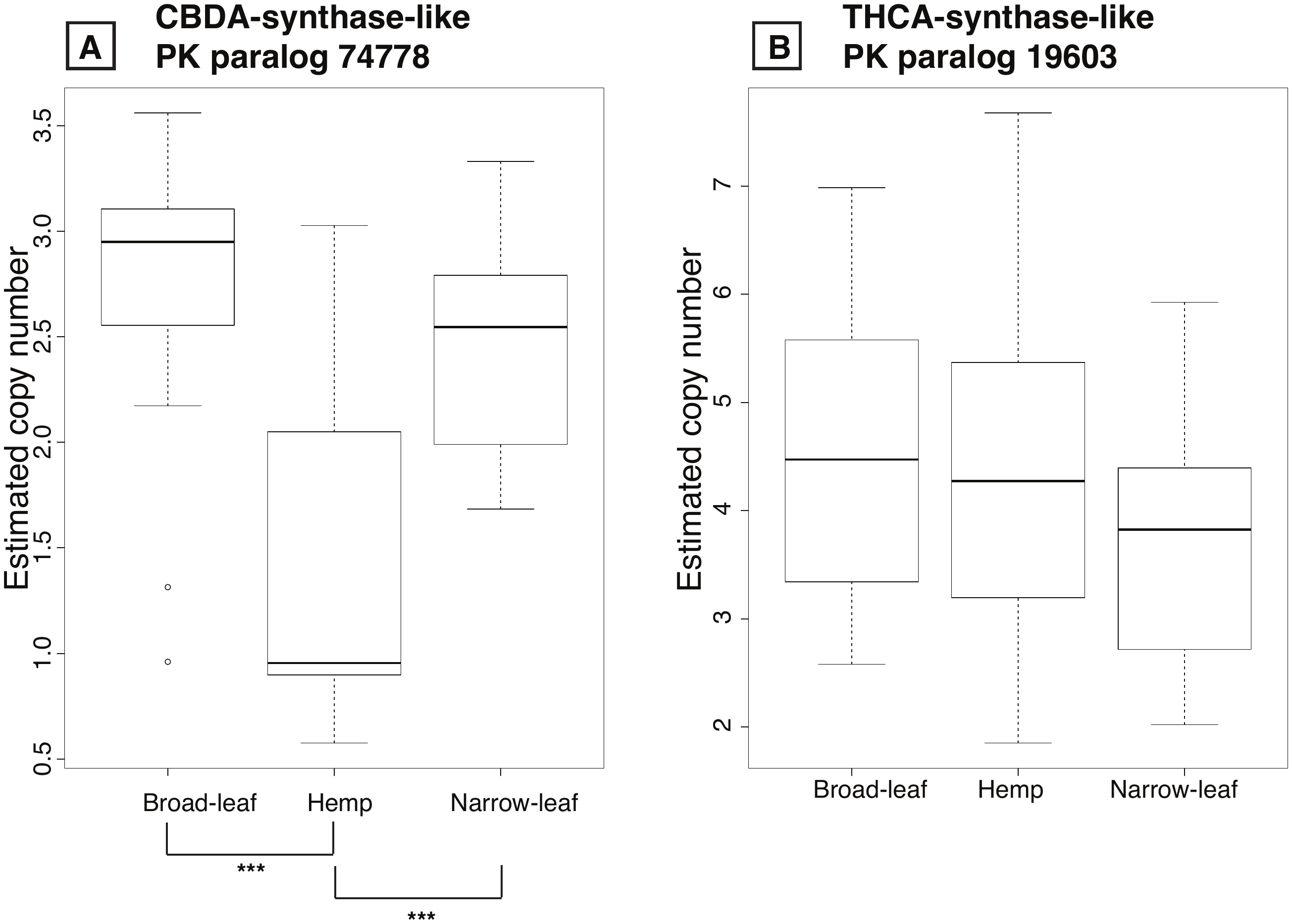

**Supporting Information Figure 1. Estimated CN by group for the two of the CBDA/THCA** **paralogs from the PK assembly.** Box plots for two of the paralogs from the 5 total paralogs of the CBDA/THCA synthase family from the PK assembly. Panel A is CBDA-like gene and panel B is the THCA-like synthase gene. Significant values between the comparisons are given in the horizontal bars below each panel: *** P<0.001, **P<0.003, *P<0.03.

Phenotypic Analysis

*CN vs chemotype correlation*

CN of THCA synthase paralog 19603 lacked significant correlation with the abundance of THCA or any other assayed cannabinoid. Of the 14 remaining genes in the analysis, only paralog 50320 from the PK assembly has a significant but negative correlation with the abundance of both CBD and CBG (Table S6). Surprisingly, CNs of the two putative olivetolic acid synthase (PK15717 and PK16618) and that of olivetolate geranyltransferase (PBBK 003891) all lack significant correlation with the abundance of any of the cannabinoids assayed.

Given that cannabinoid genes have been found physically proximal and in tandemly repeated cassettes(Weiblen et al., 2015; Grassa et al., 2018), the positive correlation between the CN of the CBDA-like paralogs in the PBBK assembly (000395 and 008242; Table S7) suggest that these paralogs could also be in close proximity and were possibly copied in tandem. Both paralogs are correlated and cluster together (Figure 2) with the PK paralog 74778 (Table S7) implying that this paralog in the PK assembly is related to CBDA production. However, the two THCA-like paralogs found in both assemblies (001774 in the PBBK assembly and 19603 in the PK assembly), which are closely related (Figure 2), do not show any correlation between them (Table S7).

After correcting for relatedness, the correlations between the levels of CBD and the CBDA-like paralog 74778 and the THCA-like paralog 19603 -both from the PK assembly-, are negative (Figure S2 panels A and B). However, the correlation between the CBDA-like paralog 74778 (Figure S2 panel A) is significant. For THC production, after correcting for relatedness, the correlations with the estimated CN of the PK paralogs 74778 (Figure S2 panel C) and 19603 (Figure S2 panel D) are both positive. Interestingly, only the correlation between THC and the CBDA-like paralog 74778 (Figure S2 panel C) is significant in contrast to the correlation between THC and the THCA-like paralog 19603 (Figure S2 panel D). In fact, paralog 19603 does not correlate with levels of any cannabinoid assayed (Table S6), possibly because we are only considering CN variation and not the effects of the specific nucleotide sequence as well as the encoded amino acid sequences, and both parameters can be important in phenotypic expression(Stranger et al., 2007). However, in one of the GLS models with THC as a response variable, both paralog 001774 and paralog 19603 were significant (t-value=3.292386, p-value= 0.0027; t-value=2.111108 p-value= 0.0438, respectively) despite both paralogs being present in both models and regardless of paralog 19603 not showing significance in the one-to-one correlation with THC (Figure S2, Table S6). This result suggests that the CN of both paralogs in each of the two assemblies influence levels of THC. Conversely, the two GLS models with CBD content as response variables did not show any significance. Still, our results and previous research suggest that CN variation may contribute to different cannabinoid phenotypes in the *C. sativa* cultivars (McKernan et al., 2015; Onofri et al., 2015).

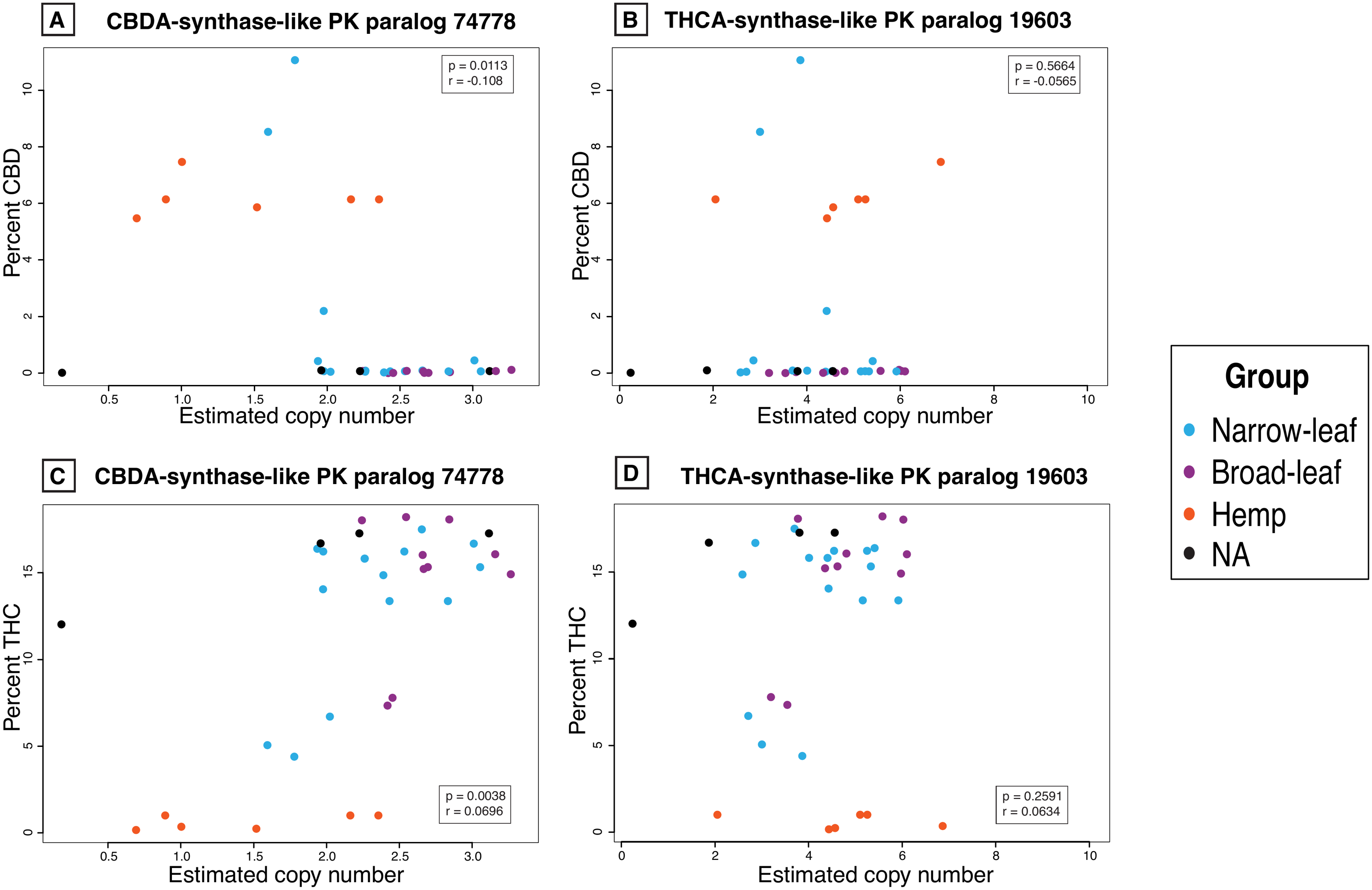

**Supporting Information Figure 2. Correlations between the percent CBD and the percent THC and the estimated CN for two CBDA/THCA synthase paralogs from the PK assembly.** The percent CBD (Panels A and B) is negatively correlated- while the percent THC (Panels C and D) is positively correlated- with CNs of both CBDA-like synthase paralog 74778 and THCA-like synthase paralog 19603 from the PK assembly. Correlation coefficient and p values in the inset after correction for relatedness. Only the CBDA-like synthase paralog 74778 is significantly correlated with both CBD (Panel A) and THC (Panel C), while the THCA-like synthase paralog 19603 lacs significance (Panels B and D). All correlation values between all genes and all cannabinoids are given in Table S6.

*The entire pathway*

Overall, both assemblies behave similarly, given that both miss genes probably due to misassembly issues. Additionally, both assemblies contain paralogs that correlate to the production of particular cannabinoids. However, we expected to see positive correlations between some of the genes and the production of particular cannabinoids, which sometimes did not happen for both assemblies. Also, the hemp group differs the most in the number of paralogs related to CBDA/THCA production despite the differences between both assemblies.

The analysis between the estimated depth of the 19 total genes (two from olivetolic acid synthase, one from olivetolate geranyltransferase and 16 from the CBDA/THCA family; Table S7) shows that CN of some of the paralogs are positively correlated independent of what gene or assembly they are related to. For example, the estimated coverage of paralog 003891 from the PBBK assembly related to the production of olivetolate geranyltransferase, is negatively correlated to the paralogs 19603 and 3498 from the CBDA/THCA synthase family from the PK assembly, and to all other paralogs from the CBDA/THCA synthase family from the PBBK assembly after correcting for relatedness. Interestingly, the two PK paralogs related to olivetolic acid synthase behave similarly after correcting for relatedness: their copy number is significantly correlated between them and each of them correlates significantly with similar paralogs (Table S7).

The difference in the estimated gene CN between assemblies could be due to the lack of a whole clade in the PK assembly (Figure 2). The presence of this clade in the PBBK assembly offers additional alignment targets for reads relative to the PK assembly, hence estimated CNs for the paralogs in the PK assembly (Figure S1) have a higher depth than those in the PBBK assembly. Similarly, reads must be shared between two copies of the CBDA paralogs in the PBBK assembly while aligning to only one in the PK assembly, again inflating the number of reads that align to paralog 74778 from the PK assembly.

Our BLAST analysis to the two newly published assemblies (Grassa et al., 2018; McKernan et al., 2018) does corroborate that in more complete assemblies cannabinoid genes from the CBDA/THCA gene family are found in multiple copies. Further research will indicate whether genes from previous steps in the pathway also have increased copies in these assemblies. Our results imply that all of these genes are found in high copies independent of the assembly used for an alignment.
